# Allele Frequency Spectrum in a Cancer Cell Population

**DOI:** 10.1101/104158

**Authors:** H. Ohtsuki, H. Innan

## Abstract

A cancer grows from a single cell, thereby constituting a large cell population. In this work, we are interested in how mutations accumulate in a cancer cell population. We provided a theoretical framework of the stochastic process in a cancer cell population and obtained near exact expressions of allele frequency spectrum or AFS (only continuous approximation is involved) from both forward and backward treatments under a simple setting; all cells undergo cell division and die at constant rates, *b* and *d*, respectively, such that the entire population grows exponentially. This setting means that once a parental cancer cell is established, in the following growth phase, all mutations are assumed to have no effect on *b* or *d* (i.e., neutral or passengers). Our theoretical results show that the difference from organismal population genetics is mainly in the coalescent time scale, and the mutation rate is defined per cell division, not per time unit (e.g., generation). Except for these two factors, the basic logic are very similar between organismal and cancer population genetics, indicating that a number of well established theories of organismal population genetics could be translated to cancer population genetics with simple modifications.

---

A tumor grows from a single cell, as has been well recognized for several decades (Muller 1950; Nowell 1976; Fidler 1978; Dexter *et al.* 1978; Merlo *et al.* 2006). Through the growth process, cells accumulate various kinds of mutations, from simple point mutations to more drastic changes at the chromosomal level, such as deletions and amplifications (Sjöblom *et al.* 2006; Wood *et al.* 2007; Network 2008; Network *et al.* 2012, 2014; Garraway and Lander 2013; Vogelstein *et al.* 2013). There are two major categories of mutations in cancer cells, driver and passenger mutations. The former are generally cell autonomous, that is, they increase the reproductive ability of the carrier cell (i.e., adaptive), while the latter have no effect on the reproductive ability (i.e., neutral). A new technology for genome sequencing from a single cell opened a new window in cancer genetics, because sequencing a number of cells from a single tumor makes it possible to identify heterogeneity in the catalog of driver and passenger mutations between cells, from which we are able to infer when and how the tumor has grown (Navin 2015).

Population genetics provides a solid theoretical framework for a wide variety of such inference methods (e.g., Nielsen and Slatkin 2013; Wakeley 2009). The coalescent (Kingman 1982; Hudson 1983; Tajima 1983) plays the central role to make the theoretical predictions of the pattern of genetic variation, which can be used to compute the likelihood of the observed variation data (Donnelly 1996; Tavaré *et al.* 1997). It concerns the history of the sampled individuals, by tracing their ancestral lineages up to the MRCA, most recent common ancestor (e.g., Nielsen and Slatkin 2013; Wakeley 2009).

One might think that the coalescent theory can be directly applied to cancer cells due to the obvious analogy; all cancer cells should follow a simple genealogy up to their MRCA. However, the direct application of the standard population genetics (i.e., organismal population genetics) to a cancer cell population may not be exactly correct because of some fundamental differences in the propagation system, as we explain below (see also Sidow and Spies 2015).

In organismal population genetics, the process can be specified by the expected number of off-springs for each individual, namely, the fitness (e.g., Crow and Kimura 1970; Ewens 1979). In the Wright-Fisher model with *N* haploids (Fisher 1930; Wright 1931), all individuals are randomly replaced every generation, and individuals with higher fitness likely produce more offsprings. In the Moran model (Moran 1962), individuals are replaced one by one, that is, one step consists of a coupling event of birth and death; one dead individual is replaced by the offspring of one randomly chosen individual from the population allowing self-replacement. Consequently, all individuals are on average replaced in *N* steps, which roughly correspond to one generation in the Wright-Fisher model. It has been well known that theoretical results under the two models are nearly identical in various cases (e.g., Crow and Kimura 1970; Ewens 1979; Wakeley 2009; Bhaskar and Song 2009). Through this random mating process either in the Wright-Fisher or Moran model, mutations that arise in the population will fix or get extinct by the joint action of random genetic drift and selection. A mutation is defined as adaptive when it increases the fitness of the carrier individual.

The evolutionary process of a cancer cell population does not follow such a simple replacement system. Figure 1 illustrates the process from cancer initiation, progression to the following rapid growth, which may be roughly divided into two major phases, and the applicability of organismal population genetics may differ depending on the phase. The first phase (Phase I) from cancer initiation to initial progression could be well handled under the organismal population genetic framework (Komarova *et al.* 2003; Iwasa *et al.* 2004; Michor *et al.* 2004). This phase is commonly modeled in a constant-size population of cells. Most theoretical models for cancer initiation suppose that a tissue consists of a number of small compartments of cells and that cancer initiation can occur in a compartment. The system starts with a normal compartment with a certain number of asexually reproducing normal cells, which is denoted by *N*_0_. *N*_0_ is usually assumed to be constant because the number of cells in a healthy tissue is maintained roughly constant by homeostatic systems, that is, cell division occurs when needed. The Moran model is more suitable to apply to this process than the Wright-Fisher model because it can be modeled such that one cell death asks for one cell division. Indeed, the Moran model has been frequently used to explore a number of problems on cancer initiation (reviewed in Michor *et al.* 2004). One of the major problems is how a cancer initiates. A compartment of a normal tissue could become a cancer when oncogenes are activated and/or tumor-suppressor genes (TSGs) are inactivated. It is believed that at least several mutational alternations in cancer genes (oncogenes and TSGs) are required for the formation of a parental cancer cell. Such accumulation of mutations in cancer genes could allow a cell to acquire typical behaviors of cancer cells, for example, avoiding apoptosis (programmed cell death) that makes it difficult to maintain the equilibrium between birth and death in the compartment, thereby shifting towards uncontrolled proliferation (neoplasia). There are a large body of theory only for the fixation process of mutations in cancer genes, especially for the inactivation of TSGs, perhaps because the problem is mathematically too simple for the activation of oncogenes (Michor *et al.* 2004). Inactivation of a TSG involves the fixation of a double-mutant, that is, both alleles have to be silenced according to Knudson’s two-hit model (Knudson 1971). This situation is very similar to the fixation process of a pair of compensatory mutations in organismal population genetics (Innan and Stephan 2001), and the results are indeed in good agreement (Iwasa *et al.* 2004). Thus, it can be considered that the applicability of organismal population genetics is quite good in Phase I because the assumption of a constant-size population roughly holds so that the stochastic process through random genetic drift works as organismal population genetics predicts.

By contrast, in the second phase (Phase II) where cells have acquired extraordinary high proliferative ability, the population grows very rapidly, and the stochastic process is less important for changing allele frequencies because most cells have very low death rates by avoiding apoptosis and their cell divisions occur independently of each other. As a consequence, a fixation of adaptive mutation hardly occurs in a cancer cell population because the spread of an adaptive mutation does not necessarily kill other cells with lower reproductive rates, as has been pointed out by Sidow and Spies (2015). This reproducing system is quite different from that organismal population genetics supposes.

We here ask how the well established theory of organismal population genetics can be applied to Phase II that presumably involves an exponential growth. In particular, we are interested in the allele frequency spectrum (AFS, or SFS: site frequency spectrum) of passenger mutations in a cancer cell population. AFS is summarized information of genotype data that are frequently used in organismal population genetics. Under the basic neutral theory of the coalescent for a constant size population (Kingman 1982; Hudson 1983; Tajima 1983) with the assumption of infinitely many sites (Kimura 1969), the expected AFS can be described in a simple form (Fu 1995), but for a non-constant size population, it is not very straightforward to obtain the expected AFS in a simple closed form. Even with any complicated demographic setting, the expected AFS can be written as a function of the expectations of coalescent times (Griffiths and Tavaré 1994, 1998), but these expectations are not easy to derive in a simple form in many cases although possible computationally (Williamson *et al.* 2005; Polanski and Kimmel 2003; Polanski *et al.* 2003). AFS provides substantial information on the past demography, making it possible to infer various demographic parameters including population size changes and migration rates (Adams and Hudson 2004; Williamson *et al.* 2005; Gutenkunst *et al.* 2009; Bhaskar *et al.* 2015; Gao and Keinan 2016).

In this article, we consider a model of a rapidly growing cancer cell population for exploring how mutations accumulate within the cancer cell population. We present some derivations for the expected AFS of derived mutations in the final tumor (at *t*_1_ in Figure 1), which could be useful to infer when the exponential growth started and how fast the tumor has grown. There has been extensive works on a cancer cell population by Durrett (Durrett 2013, 2015), who provided approximate formulas to the sample-based AFS. We have here obtained analytical expressions of the expected AFS in a near exact form (only continuous approximation is involved) by both forward (branching theory) and backward (coalescent theory) treatments. The former is in a simpler form that is useful for intuitive understanding of the process, while the latter provides a solid theoretical framework for coalescent (backward) simulations of a cancer cell population. Our near exact result is compared with Durrett’s approximate formulas, together with some simulation results.

It should be noted that our interest is in passenger mutations in the second phase with the assumption of no driver mutations so that the increase of the cancer cell population size can be approximated by an exponential function. There is no doubt that a number of driver mutations are involved in the first phase (e.g., Knudson 1971; Michor *et al.* 2004; Sjöblom *et al.* 2006; Network *et al.* 2014), but there are extensive debates on the potential role of driver mutations in the second phase. Some authors suggest that the role of driver mutations may be quite limited after the original cancer cell is stablished and most mutations occurs in the following growth phase may be passengers (Uchi *et al.* 2016; Sottoriva *et al.* 2015), whereas some point out the importance of driver mutations (Williams *et al.* 2016; Waclaw *et al.* 2015; Marusyk *et al.* 2014). Because our model assumes no driver mutations in the second growth phase, the theoretical result could be used as a null model for testing the role of driver mutations in the second phase.

**Figure 1.**
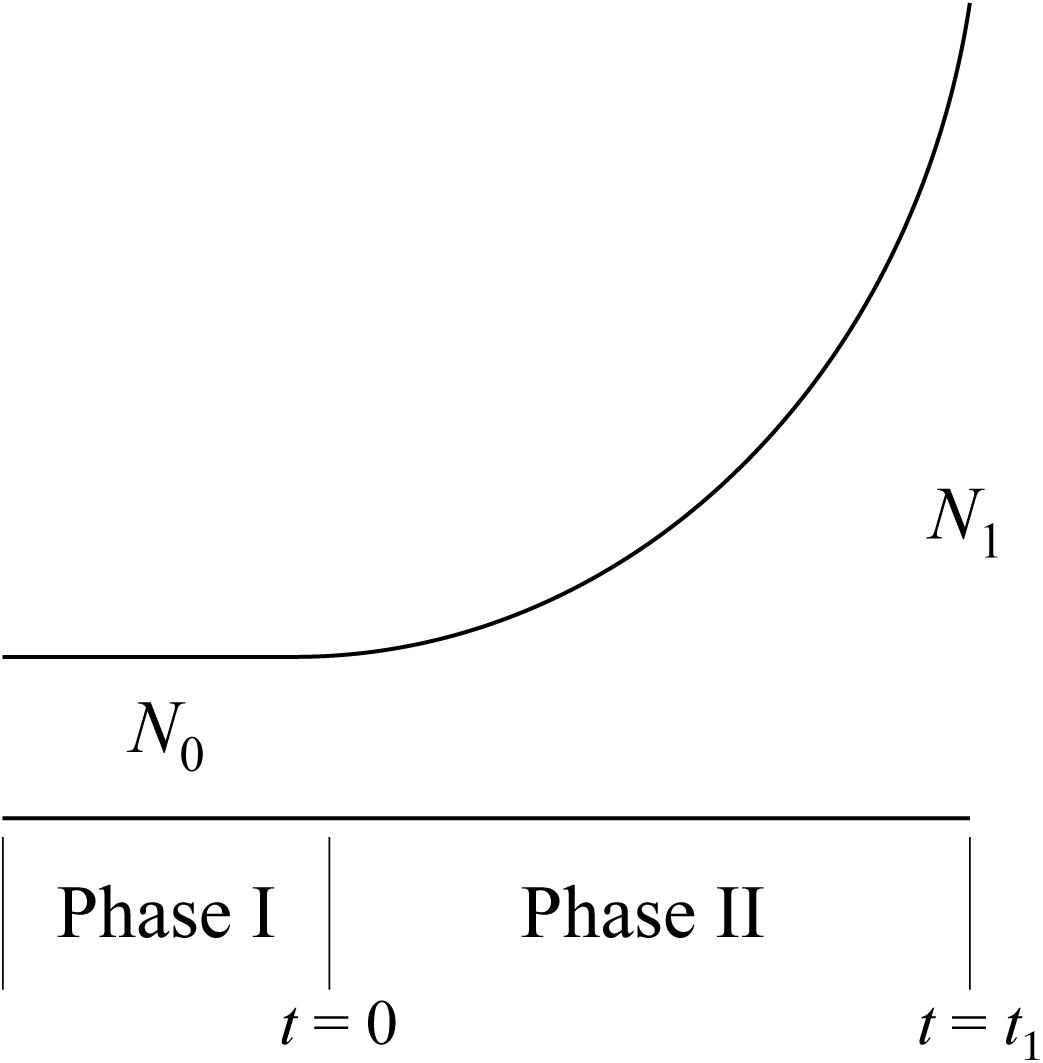
Illustrating the model of the growth of a cancer cell population.

## MODEL

Our model (Figure 1) considers an exponentially growing population starting with *N*_0_ asexually reproductive cells. The reproductive ability of a cell is specified by the cell division rate (birth rate) and death rate per time unit, denoted by *b* and *d*, respectively, which are assumed to be constant over time. The tumor starts growing at time *t* = 0, and let *N*(*t*) be the number of cells at time *t*. For convenience, we define *t*_1_ such that *N*(*t*_1_) = *N*_1_ is satisfied for the first time. Under this setting, because it is obvious that the Moran model does not work, we use the branching process.

We assume *b* ≫ *d* so that the tumor grows approximately exponentially at rate *r* = *b* − *d* and the number of cells at *t* is approximately given by

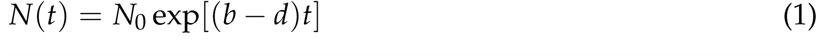

This equation is a very good approximation unless *N*_0_ is very small. Note that in reality *N*(*t*) follows some distribution, but our deterministic treatment on *N*(*t*) does not affect the following results much.

The rate of passenger mutation is given such that at each cell division one of the daughter cells receives a novel mutation at rate *μ*. We assume a very small rate per site so that the assumption of the infinite-site model (Kimura 1969) holds.

### Forward Treatment by Branching Process

We aim to obtain the expected derived allele frequency spectrum (AFS) when the total number of cells is *N*_1_ (i.e., *t* = *t*_1_), where we assume that *N*_1_ ≫ *N*_0_. The expected number of passenger mutations that are shared by *i* cells at time *t* = *t*_1_ is denoted by *S*(*i*, *μ*, *t*_1_). Because of our deterministic assumption (i.e, Equation (1)), *t*_1_ is given such that it satisfies *N*_1_/*N*_0_ = exp[(*b* − *d*)*t*_1_].

We first consider how many cells at *t* = *t*_1_ share a particular mutation that occurred at *t* = *t*_1_ – *t′*. We here use the well-known formula under the branching process: the probability density function (pdf) of the number of daughter cells (*i*) of a particular single individual after *t′* time units is given by:

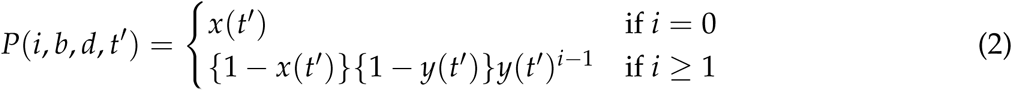

(Bailey 1964), where

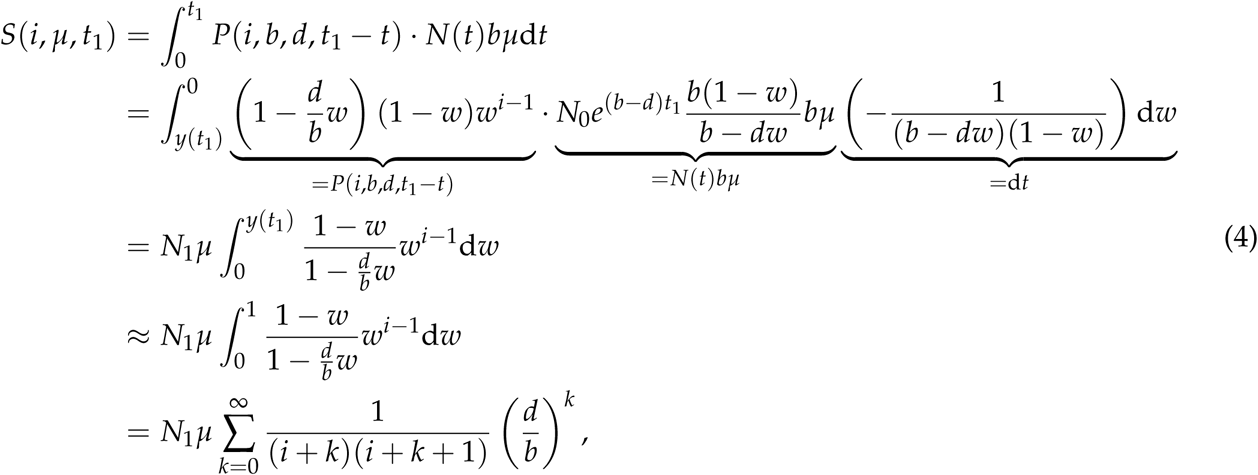

This formula provides an unconditional distribution of the number of individuals having a specific origin, which is independent of the total population size. Nevertheless, we use this formula by ignoring the effect of the total population size. This simplification is reasonable and the effect on the theoretical treatments is negligible even though it is technically possible that *i* exceeds the total population size. This is because *i* is usually not a large number unless *N*(*t*_1_ − *t′*) is unrealistically small.

We then obtain *S*(*i*, *μ*, *t*_1_), the expected number of mutations with frequency *i* in the final tumor by considering all potential mutations that occur 0 < *t* < *t*_1_. Because the population mutation rate at time *t* is *N*(*t*)*bμ*, we obtain *S*(*i*, *μ*, *t*_1_) for *i* ≥ 1:

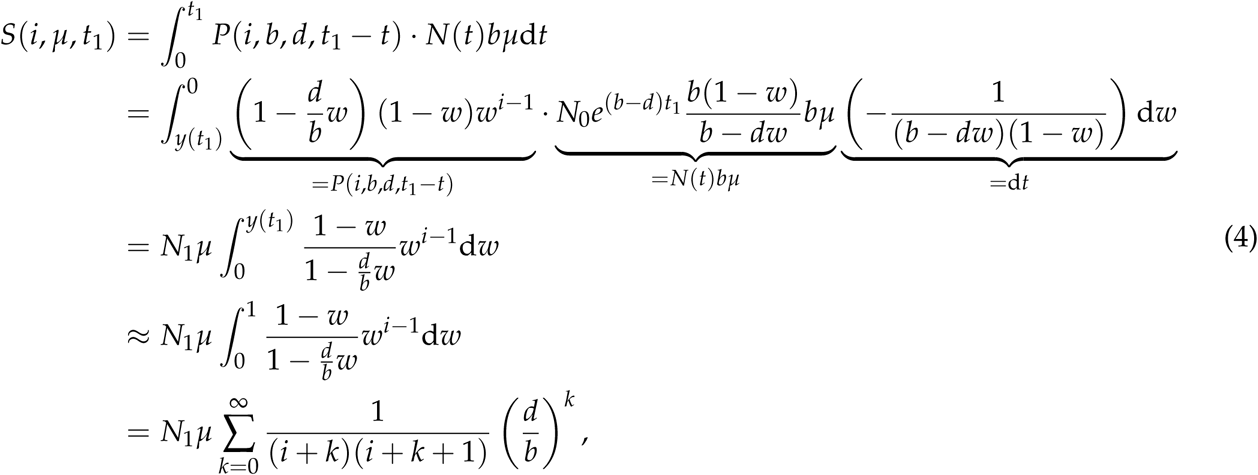

where we set *w* = *y*(*t*_1_ – *t*) and assume *y*(*t*_1_) ≈ 1 and *N*_0_ is very small. We again note that because of the nature of our approximation, it is possible to compute *S*(*i*, *μ*, *t*_1_) even for i > *N*_1_. For a practical calculation of *S*(*i*, *μ*, *t*_1_), however, this treatment should not matter so much as mentioned above. Equation (4) means that the relative frequency distribution of *S*(*i*, *μ*, *t*_1_) is determined by the ratio of *d* to *b*, while *N*_1_*μ* determines the absolute number of mutations.

It is straightforward to obtain the expected normalized AFS (pdf of *i* given a segregating mutation, i.e., *i* = (1, 2, 3,…, *N*_1_)) as

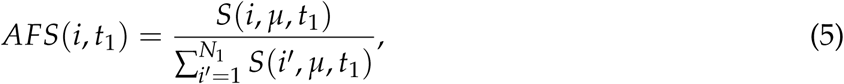

where, for a large *N*_1_, the denominator of eq.(5) is approximated by

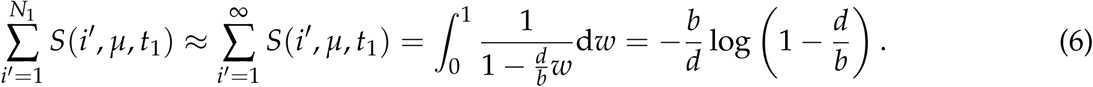

Of particular importance is the case of *b* ≫ *d*, that is, the population grows very rapidly, where Equation (4) becomes

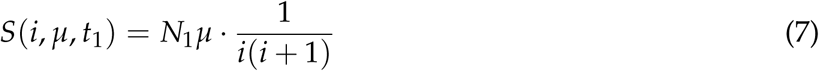

and

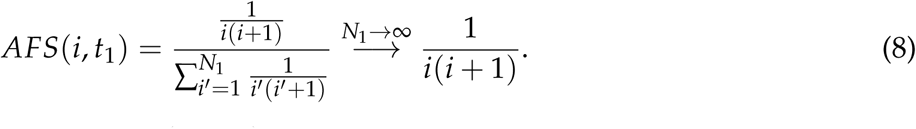

At this limit, it is interesting to note that *S*(*i*, *μ*, *t*_1_) is independent of *b* or *d*.

We can consider the opposite extreme, *b* ~ *d*, where the underlying assumption of our calculation (*i.e*., the population grows exponentially) is obviously broken. Nevertheless, if we formally proceed our calculation by taking the limit *b* → *d*, we obtain

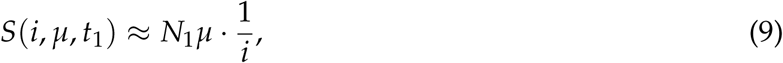

which reproduces the result for a Moran process in a constant-size population (Fu 1995; Griffiths and Tavaré 1998; Wakeley 2009). This is not a coincidence because our assumption *b* = *d* with deterministic treatment simply means a constant size population, but this equation does not work well in our randomly reproductive population without keeping the population size constant. For AFS, we have

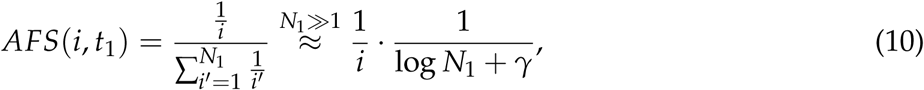

where *γ* ≡ 0.577215… is the Euler’s constant.

We performed forward simulation to check how our equations work. Our simulations assumed that *N*_0_ = 10, *N*_1_ = 10^5^, *μ* = 10^−4^, *b* = 4, and *d* = {0, 0.2, 0.4, 1}, and Figure 2 shows the average spectra (up to *i* = 25) over 10^5^ simulation runs. Theoretical results based on Equation (4) are shown in closed circles. It is demonstrated that Equation (4) is in excellent agreement with the simulation results (colored open boxed) for all four cases. We further compare the values computed by Equation 7 (filled triangles in Figure 2), which is a simple approximation to Equation (4) when *b* ≫ *d*. We find that Equations (4) and (7) produce almost identical numerical values, which are indistinguishable in Figure 2, indicating that the simple approximation works very well when *d* = 0. Furthermore, Equation (7) could be in fairly good agreement with the results of Equation (4) with *d*/*b* = 0.05, indicating that the simple form (Equation (7)) can be a good approximation when *d*/*b* ≪ 0.05.

For applying our theoretical result to data, it is more convenient to consider a sample rather than the entire cell population. Suppose that n random cells are sampled from the population. Then, the expected number of mutations that are shared by *i* (1 ≤ *i* ≤ *n*) cells in a sample of size *n* is given by

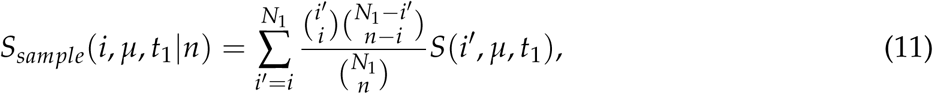

which is, for a large *N*_1_, approximated by using a Poisson distribution as

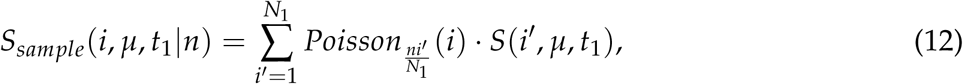

**Figure 2.**
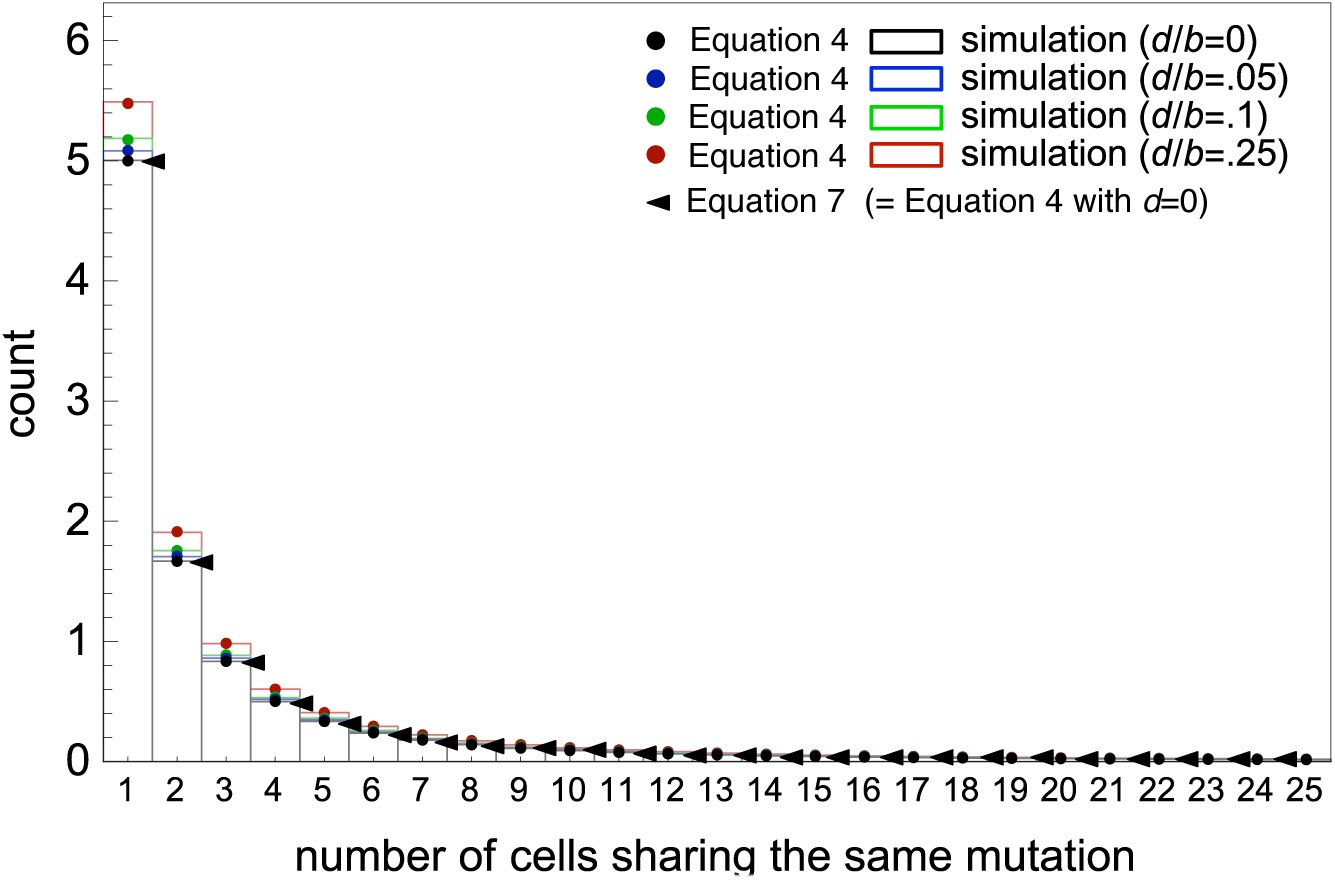
Population allele frequency spectra, *AFS*(*i*, *t*_1_), when *d*/*b* = {0, 0.05, 0.2, 0.25}. The theoretical results from Equations (4) and (7) are compared with simulations. Forward simulations were performed with *N*_0_ = 10, *N*_1_ = 10^5^, *μ* = 10^−4^, *b* = 4, and *d* = {0, 0.2, 0.4, 1}. It should be noted that Equation (4) with *d* = 0 is identical to Equation (7).

where

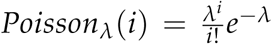
 Then, it is straightforward to obtain normalized sample AFS. If we include fixed mutations, the normalized sample AFS is given by

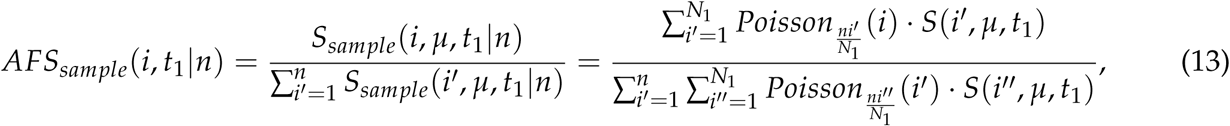

and if fixed mutations are ignored

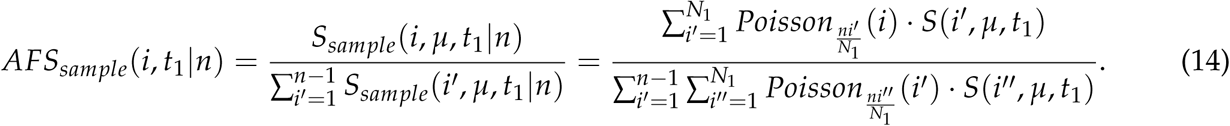

### Backward Treatment by the Coalescent

The coalescent is one of the major theories in organismal population genetics. It is a sample-based theory: The lineages of sampled individuals are traced backward in time until they coalesce into their MRCA (most recent common ancestor). We here apply this logic to a sampled cells from a tumor, and obtain essentially the same theoretical results as those from the forward treatment (i.e., Equations (11 – 14)).

Let us consider a pair of random (different) cells from the final tumor with *N* cells, where *N* is already a large number. Because the following argument works at any time in Phase II (assuming *N*_0_ is very small), we shall use *N* for the population size rather than *N*_1_. We consider backward time *τ* from the present, such that the present time is set to *τ* = 0. Let *T*_2_ be the time it takes for the two lineages to coalesce. We consider an infinitesimally small time interval Δ*τ* such that at most one event (birth or death) can occur. The conditional probability that the population size was *N* – 1 at time Δ*τ* backward, conditioned on that the present population size is *N* is given by

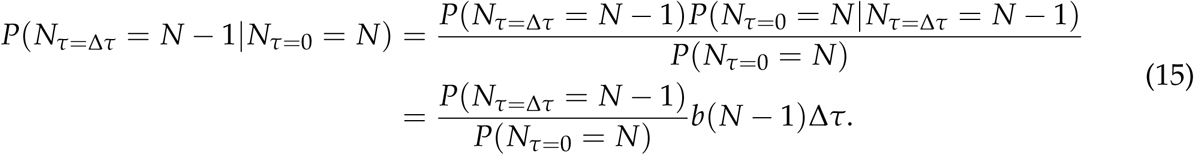

The probabilities *P*(*N*_Δ*τ*_ = *N* – 1) and *P*(*N*_0_ = *N*) can be calculated based on the forward process, but for a large *N* it is expected that their difference is at most of order Δ*τ*, so the leading term of the expression above is *b*(*N* – 1)Δ*τ*. This equation represents the probability that a birth event occurred in the interval Δ*τ*. The birth event can cause the coalescence between two specific lineages, with probability

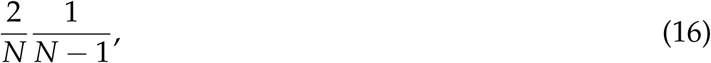

and therefore the probability of coalescence is, up to the first order of Δ*τ*, given by

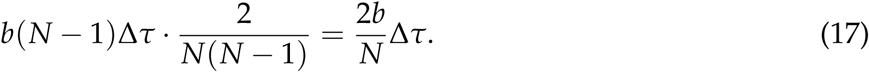

In the mean time, we must take into account the fact that the population is shrinking at rate *r* = *b* – *d* backward in time (Slatkin and Hudson 1991). The rate of coalescence between two lineages at time *τ* is approximated by

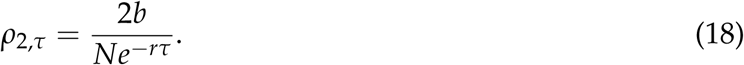

Note that this formula is consistent with a well-known formula for the Moran process when the population size is fixed (e.g., Wakeley 2009), namely, setting *b* = *d* = 1 reproduces

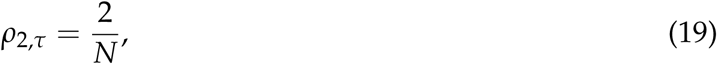

which is the per-generation rate of coalescence for the Moran model.

Let *P*_2_(*τ*) be the probability that the coalescence between the two lineages have not occurred yet by time *τ*, for which the following differential equation holds:

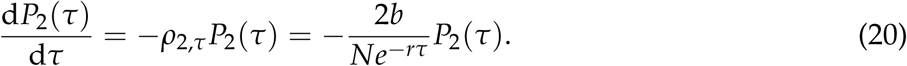

With *P*_2_(0) = 1 as a boundary condition, the solution is given by a double exponential function:

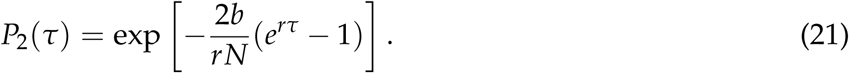

Therefore, the density function of coalescent time *T*_2_ is given by

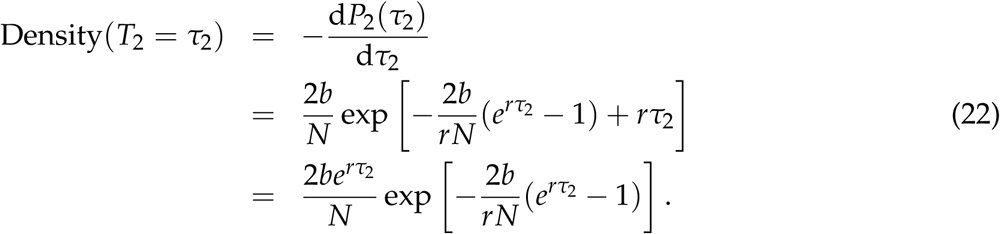

Following the same logic, for *k* (> 2) cells, we have

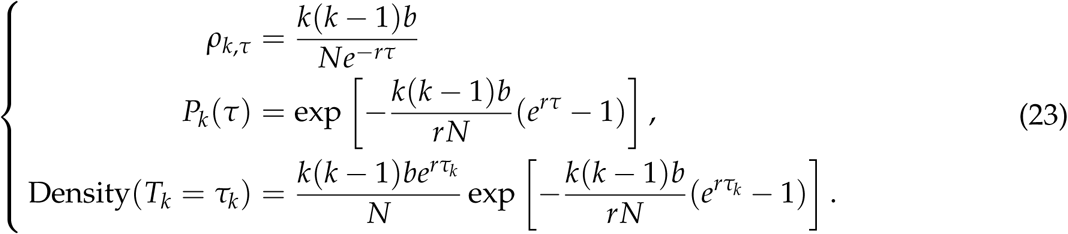

In order to consider the coalescent process of *n* sampled cells up to their MRCA, we are interested in the joint pdf of {*T*_2_, *T*_3_,…, *T*_*n*−1_, *T_n_*}, which is given by

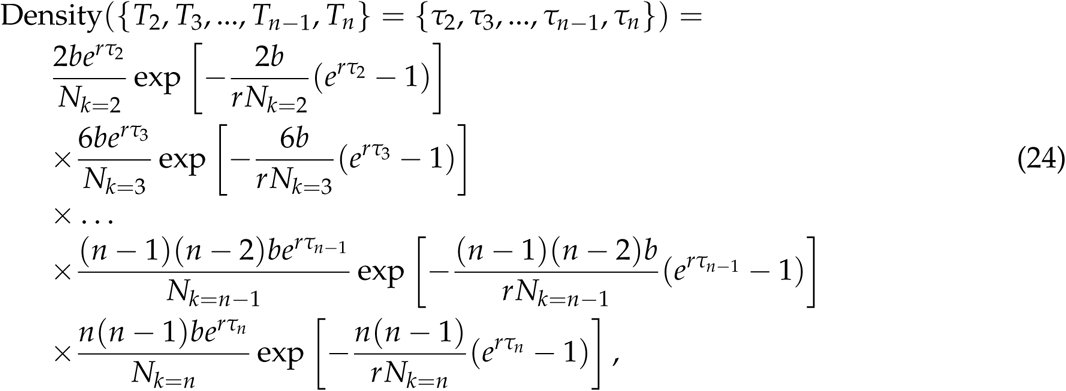

where *N*_*k*=*j*_ is the population size at the moment when the original *n* lineages coalesce up to *j* lineages. In other words, *N*_*k*=*j*_ is the population size 
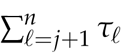
 time units before the present. Thus, the coalescent times are not independent one another, that is, *T_j_* is given conditional on 
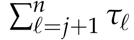
.

We can generate a (*n* − 1)-turple of coalescent time, {*τ*_2_, *τ*_3_,…, *τ*_*n*−1_, *τ*_n_}, from the joint distribution (24) in the following way. First we set *N*_*k*=*n*_ = *N*_1_, that is the size of the population where *n* samples are originally taken. Then, generate a random number *τ_n_* according to the density distribution given by (23). Next, set *N*_*k*=*n*−1_ = *N*_1_ exp[−*rτ_n_*], and generate a random number *τ*_*n*−1_ according to the density distribution given by (23). The value of *N*_*k*=*n*−2_ is then set to *N*_*k*=*n*−2_ = *N*_1_ exp[−*r*(*τ_n_* + *τ*_*n*−1_)] and *τ*_*n*−2_ is generated, and so on.

The expected normalized AFS under this coalescent process can be described as

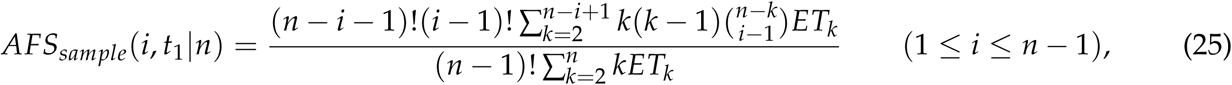

where *ET_k_* is the expectation of *T_k_* that can be obtained from (24) (Griffiths and Tavaré 1998; Wakeley 2009). For the absolute number of mutations that exactly *i* individuals in a sample of size *n* have, *S*_*sample*_(*i*, *μ*, *t*_1_ | *n*), it is not difficult to see that

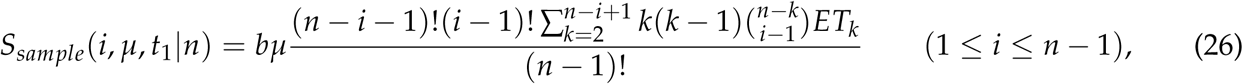

holds. This is because *ET_n_* contributes to *S*_*sample*_(1, *μ*, *t*_1_ | *n*) in the form of 2*b* · *μ* · (1/2) · *nET_n_* = *bμnET_n_*, where 2*b* is the backward rate of birth event per lineage, *μ* is the mutation rate, (1/2) is the chance that the focal lineage receives a mutation at a single birth event, *n* is the total number of independent lineages, and *ET_n_* is the expected duration during which there are *n* independent lineages. As the expected coalescent time *ET_k_* can be computed based on the numerical procedure provided above, it is straightforward to numerically calculate the sample AFS with Equations (25) and (26).

**Figure 3.**
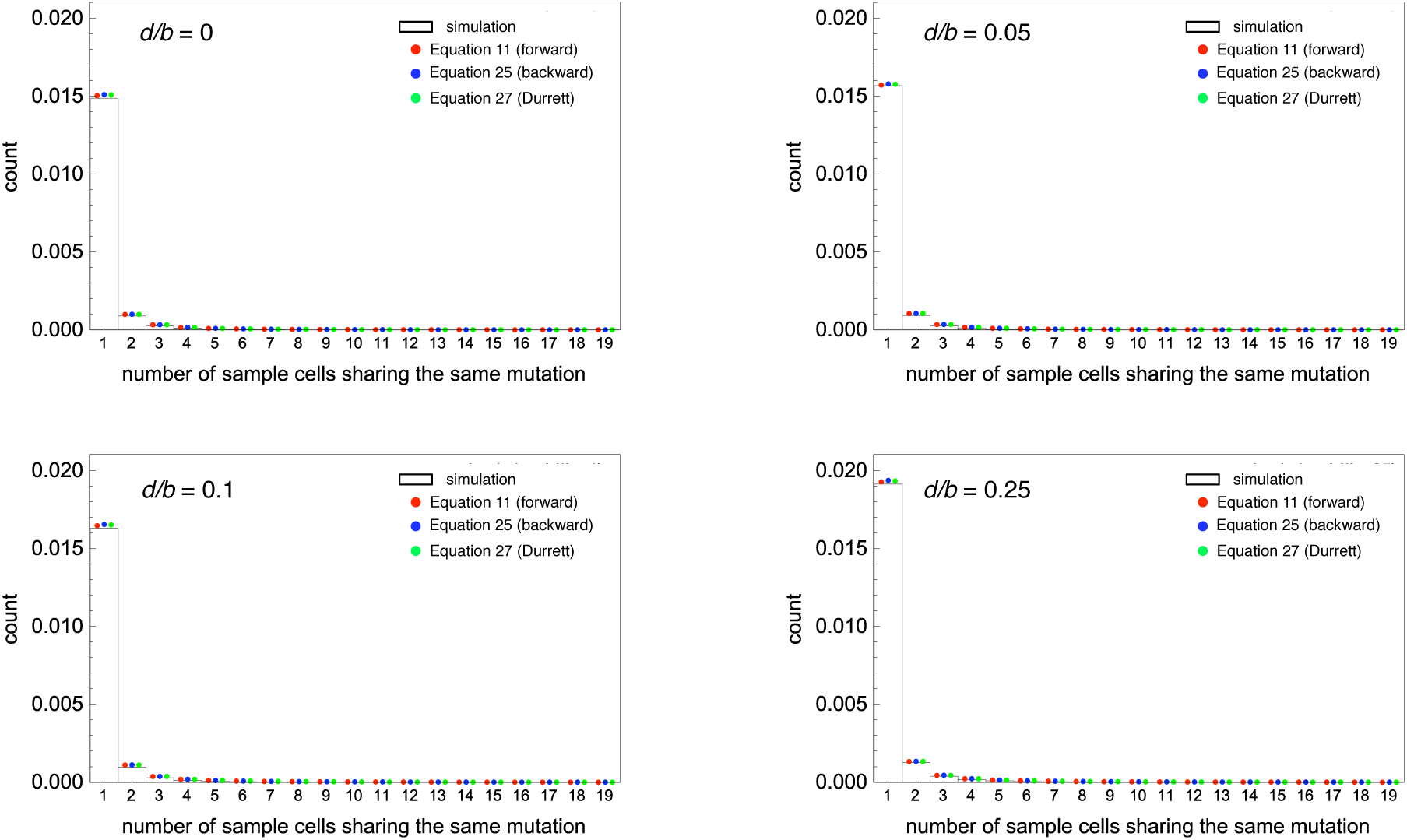
Population allele frequency spectra, *AFS*(*i*, *t*_1_), when *d*/*b* = {0, 0.05, 0.2, 0.25}. The theoretical results from our forward and backward treatments and Durrett’s approximation (28) are compared with simulations. The simulation results are icentical to those used in Figure 2.

## DISCUSSION

This article considers a model of a rapidly growing cancer cell population for exploring how mutations accumulate within the population. The expected AFS of derived mutations is obtained in a near exact form by both forward (branching theory) and backward (coalescent theory) treatments. Durrett (2013, 2015) obtained two approximate formulas to the sample-based AFS:

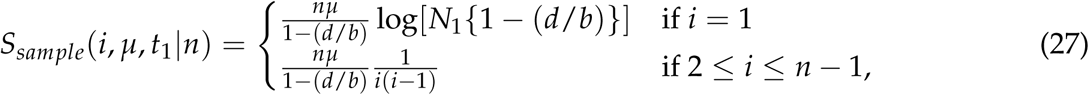

which was ultimately improved to be

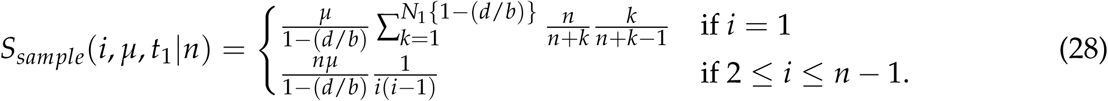

In Figure 3, the numerical results from our forward and backward derivations (i.e., Equations (11) and Equation (26) are compared with Durrett’s two approximations (Equations (27) and (28)). There is nothing surprising that Equations (11) and (26) are in excellent agreement because they are in near exact forms. In addition, we find Durrett’s improved approximation (28) is extremely good, while the first approximation (27) would overestimate the singleton frequency (not shown).

The advantage of our near exact expressions over Durret’s great approximation is that our theory would provide some mathematical intuitions, which could be useful for data analysis. (i) First, our forward expression (4) can be approximated to a very simple form for a large *r* = *b* − *d*:

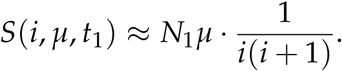

This means that *N*_1_*μ* determines the absolute number of mutations and the relative frequency is converged to 
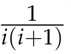
 with *r* → ∞, which is independent of the growth rate. Provided that the growth rate of a typical cancer cell population is very large, AFS may not be very informative to estimate the growth rate. Rather, *N*_1_*μ* may be more informative biologically because the mutation rate (*μ*) may be easily estimated if *N*_1_ is given. It may not be very difficult to obtain a rough estimate of *N*_1_ from the size of tumor. One might think that this implication seems odd: What if a tumor has grown from *N*_0_ = 1 to *N*_1_ = 10^10^ in an hour or so? An hour could be too short to accumulate mutations. To address this question, we should note that how short the time is taken, it has to have involved at least *N*_1_ − *N*_0_ cell divisions and that the mutation rate is defined per cell division, not per time unit.
(ii) Second, our backward expression for the coalescent time is

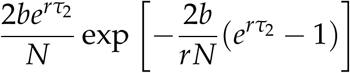

(identical to Equation (22)), which is in a similar form to that under the standard coalescent in organismal population genetics:

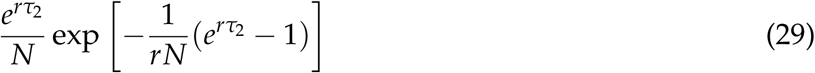

(Slatkin and Hudson 1991). The difference between those two expressions can easily be explained; the factor 2 in the former equation reflects the fact that the our model assumes overlapping generation, while Slatkin and Hudson (1991) did not (e.g., Wakeley 2009). After neglecting this factor 2, these two equations are completely equivalent when the birth rate of a cell is *b* = 1 in our model. By comparing these two equations, the expression of Slatkin and Hudson (1991) for organismal population genetics is a special case of our expression. In other words, the well-established backward theory of organismal population genetics can be directly used to a cancer cell population by introducing a scale factor *b* that determines the relative rate of coalescent and population shrinkage (in backward).

One may think from our formulas of coalescent time (22) that the absolute values of *b* and *r* = *b* − *d* jointly specifies the process. This is indeed true if we are interested in the absolute length of waiting time until coalescence. On one hand, if only allele frequency spectrum is of interest, those absolute values are much less important. Rather, the ratio of *d* to *b*, namely *d*/*b*, is a crucial determinant of the spectrum, as is obvious in Equation (4), which explicitly tells us that it is the case because it depends on *b* and *d* only through *d*/*b*. It may be difficult to see this fact in our backward formula (e.g., (22)), but if we rescale backward time and introduce a new timescale *τ′* by *τ′* = *bτ*, then Equation (20), for example, changes to

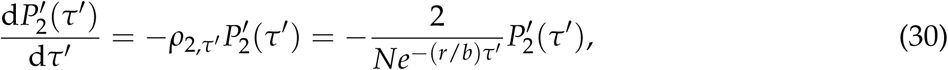

which depends on b and d only through 
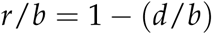
 and therefore only through *d*/*b*. This intuitively makes sense because in our cancer model, mutation occurs only at birth events, so the absolute waiting time until a birth event occurs is irrelevant when we focus on AFS of a population/sample.

(iii) Third, our expressions are based on solid derivations. Therefore, the basic logic behind our derivations can be applied to more complex growth pattern as long as *b* ≫ *d*. It can be considered that the growth of a cancer cell population may not be necessarily exponential. Driver mutations could increase the growth rate, while the growth process may slow down if the availability of resources such as space, oxygen, and other nutrients is limited. Such change of the growth curve may be incorporated by replacing Equation (1), which will be involved in the integration in (4) in the forward treatment and in the rate of coalescent specified by (18) in the backward treatment.

In summary, assuming an exponentially growing cancer cell population, we obtained near exact expressions of AFS (only continuous approximation is involved) from both forward and backward treatments. The former is in a simpler form and enhance our intuitive understanding of the process, while the latter provides a theoretical framework for coalescent (backward) simulations of a cancer cell population. Our theoretical results show that the difference from organismal population genetics is mainly in the coalescent time scale and the mutation rate is defined per cell division, not per time unit (e.g., generation). Except for these two factors, the basic logic are very similar between organismal and cancer population genetics. Therefore, a number of well established theories of organismal population genetics could be translated to cancer population genetics with simple modifications.

## Acknowledgements

This work is in part supported by grants from the Japan Society for the Promotion of Science (JSPS) to HO and HI.

